## Supplementary Figs 1 and 2 for "A myosin II-based nanomachine devised for the study of Ca^2+^-dependent mechanisms of muscle regulation"

### Supplementary Materials

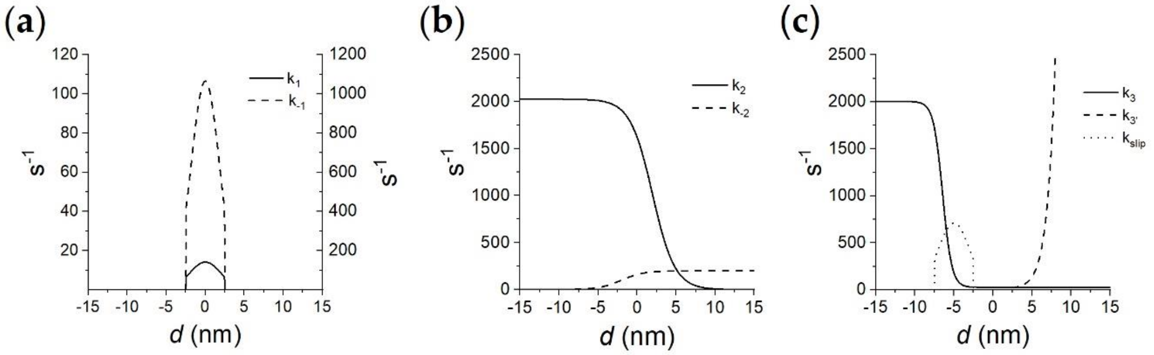


**Supplementary Fig. 1.** Dependence of the rate functions for the state transitions on *d*. Where the transition is reversible, ki is the forward and k-i is the backward rate constant. **(a)** step 1, attachment reaction, k1 continuous line, k-1 dashed line. **(b)** Step 2, force generating transition, k2 continuous line, k-2 dashed line. **(c)** Step 3, detachment reaction, k3 continuous line. The slipping of A2 to the next actin farther from the centre of the sarcomere is governed by $k_{slip}$ (dotted line). Detachment from an attached state following slipping, $k_{3'}$ dashed line.

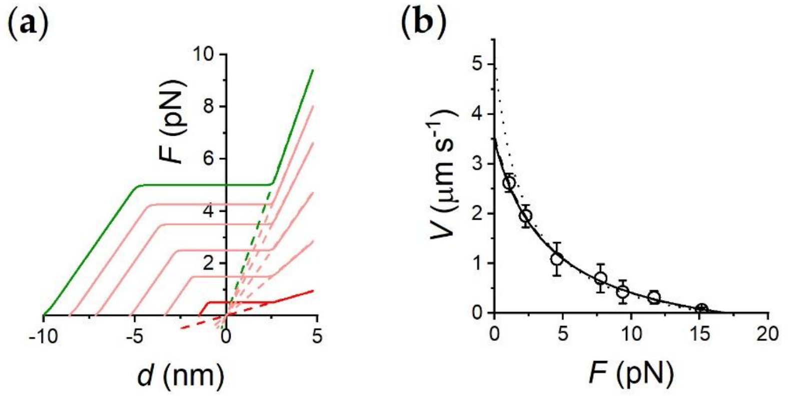


**Supplementary Fig. 2.** (**a**) Force profile of the attached states of the motor as a function of *d*, the relative position between the motor and the actin monomer. A1 state, dashed line and A2 state, continuous line. Green, force profile of correctly oriented motors; red, of motors 180° away from the correct orientation; pink, four profiles corresponding to four intermediate motor orientations. (**b**) *F*-*V* relation simulated for a number of available heads $N$ = 16 with the model in Ref. [10] (dotted line) and with the implemented version that takes into account the effect of the random orientation of motors on both force and the step size (continuous line). Experimental data (circles) and their fit with Hill equation (dashed line) from Fig. 2d.
